# ChIA-DropBox: a novel analysis and visualization pipeline for multiplex chromatin interactions

**DOI:** 10.1101/613034

**Authors:** Simon Zhongyuan Tian, Daniel Capurso, Minji Kim, Byoungkoo Lee, Meizhen Zheng, Yijun Ruan

## Abstract

ChIA-Drop is a new experimental method for mapping multiplex chromatin interactions with single-molecule precision by barcoding chromatin complexes inside microfluidics droplets, followed by pooled DNA sequencing. The chromatin DNA reads with the same droplet-specific barcodes are inferred to be derived from the same chromatin interaction complex. Here, we describe an integrated computational pipeline, named ChIA-DropBox, that is specifically designed for reconstructing chromatin reads in each droplet and refining multiplex chromatin complexes from raw ChIA-Drop sequencing reads, and then visualizing the results. First, ChIA-DropBox maps and filters sequencing reads, and then reconstructs the “chromatin droplets” by parsing the barcode sequences and grouping together chromatin reads with the same barcode. Based on the concept of chromosome territories that most chromatin interactions take place within the same chromosome, potential mixing up of chromatin complexes derived from different chromosomes could be readily identified and separated. Accordingly, ChIA-DropBox refines these “chromatin droplets” into purely intra-chromosomal chromatin complexes, ready for downstream analysis. For visualization, ChIA-DropBox converts the ChIA-Drop data to pairwise format and automatically generates input files for viewing 2D contact maps in Juicebox and viewing loops in BASIC Browser. Finally, ChIA-DropBox introduces a new browser, named ChIA-View, for interactive visualization of multiplex chromatin interactions.

## Introduction

Over the past decade, experimental methods have been developed to map the three-dimensional (3D) organization of genomes. These methods detect bulk chromatin interactions, genome-wide chromosome conformation capture (Hi-C) ^1^ or enrich for chromatin interactions involving a specific protein, chromatin interaction analysis with paired-end tag sequencing (ChIA-PET) ^2^. ChIA-PET and Hi-C have been extensively applied and have suggested complex chromosomal folding structures. However, these methods are based on nuclear proximity ligation, and reveal only pairwise contacts at population-level aggregated from millions of cells. Thus, the true multiplex nature of chromatin complexes at the single-molecule level had remained unexplored. A new direction is to develop single molecule approaches to directly detect multiplex chromatin interactions from either single cells ^3^ or bulk cells ^4^.

Recently, we developed ChIA-Drop, a novel experimental method for detecting multiplex chromatin interactions with single-molecule precision via droplet-based and barcode-linked sequencing ^5^. In ChIA-Drop, a crosslinked and fragmented chromatin sample is directly loaded into a microfluidics system (10X Genomics). The chromatin sample can either be without specific enrichment (similar as for Hi-C) or enriched for a specific protein factor by immunoprecipitation (similar as for ChIA-PET). Once the sample is loaded in the microfluidics system, the chromatin complexes are partitioned into Gel-bead-in-Emulsion (GEM) droplets. Each droplet then contains a gel bead of unique DNA oligonucleotides and reagents for linear amplification and barcoding of the chromatin DNA templates (Fig. 1a). The barcoded amplicons with GEM-specific indices can then be pooled for standard high-throughput sequencing, and the single-molecule chromatin complexes can be inferred computationally through processing and analysis of the sequencing data.

**Figure 1:**
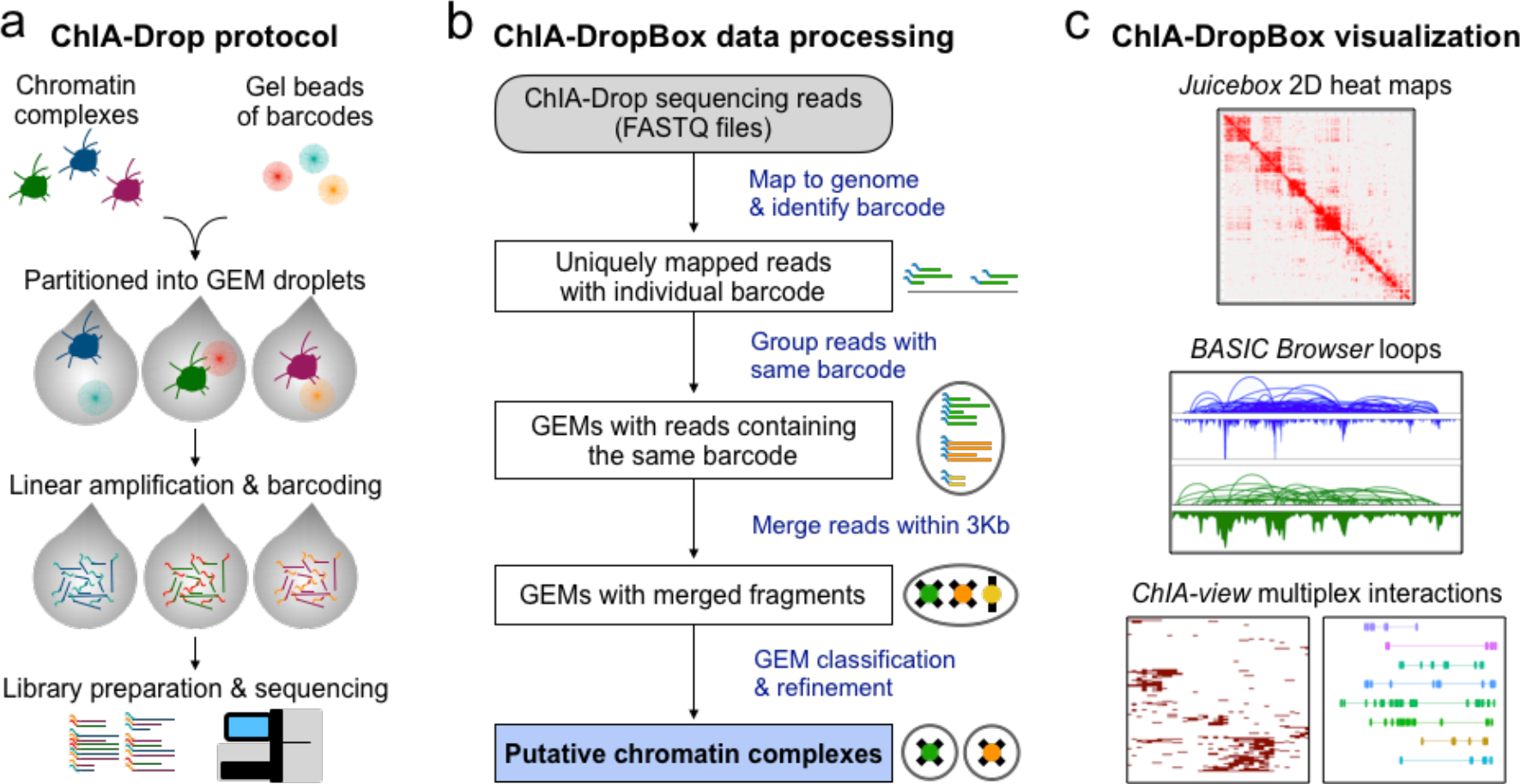
ChIA-Drop protocol and ChIA-DropBox computational pipeline. (**a**) Schematic of the ChIA-Drop experimental protocol. A dual-cross-linked and fragmented sample of chromatin complexes is loaded directly into the 10X Genomics microfluidic device. The sample can be protein enriched (immunoprecipitated) or unenriched. In the microfluidics device, chromatin complexes are partitioned into gel-bead-in-emulsion (GEM) droplets for barcoding and linear amplification. Then the barcoded amplicons are pooled for standard high-throughput sequencing. (**b**) Schematic of the ChIA-DropBox computational pipeline. First, the 10X Genomics Chromium software suite is applied to align reads to the reference genome and to identify the GEM barcode of each read. ChIA-DropBox then groups uniquely mapped reads with the same barcode to reconstruct the GEMs. Within each GEM, reads that overlap in their linear genomic alignments within 3Kb are merged into chromatin fragments (considered with the fragment size from library preparation). Finally, considering the concept of chromosome territories, the GEMs are refined into purely intra-chromosomal putative chromatin complexes. (**c**) ChIA-DropBox enables high-resolution data visualization via several approaches. First, ChIA-DropBox converts the data to pairwise format and generates input files for existing visualization tools: 2D contact map views via Juicebox and loop views via BASIC Browser or other genome browsers. Importantly, ChIA-DropBox also introduces its own custom visualization tool specifically for viewing multiplex chromatin complexes, called the “ChIA-View”, which includes a cluster view and a fragment view, implemented by *R* Shiny.

We have successfully generated dozens of ChIA-Drop (no enrichment) and RNAPII-enriched ChIA-Drop libraries, and have demonstrated that the method is simple, robust, and reproducible for detecting multiplex chromatin interactions ^5^. The ChIA-Drop experimental method has many advantages: it can analyze chromatin samples directly without the need to purify chromatin DNA, it is independent of proximity ligation, it detects multiplex chromatin interactions with single-molecule precision, it is simple and robust to perform experimentally, and it uses standard pooled sequencing of the barcoded fragments. However, the pooled sequencing means that computational tool is essential to obtain interpretable data by reconstructing and refining the multiplex chromatin complexes. Thus, during the course of ChIA-Drop development, we also developed a concomitant computational pipeline ChIA-DropBox with an advanced and open-source framework to tackle the major challenge of going from raw ChIA-Drop sequencing reads for mapping, analyzing, and visualizing multiplex chromatin interactions. Here, we describe the modules in the ChIA-DropBox pipeline, a novel analysis and visualization pipeline for multiplex ChIA-Drop data (Fig. 1, Supplementary Fig. 1).

## Results

### Overview of ChIA-DropBox

The ChIA-DropBox pipeline is composed of four modules, mapping of sequencing reads, assembling of chromatin fragments and chromatin droplets, refinement of chromatin complexes, and data visualization. Briefly, the 10X Genomics Chromium software suite pipeline is first applied to align reads to the reference genome and to identify the GEM barcode of each read. ChIA-DropBox then groups uniquely mapped reads with the same barcode to reconstruct the GEMs. For each GEM, reads with overlapping genomic coordinates within 3 Kb are merged into “chromatin fragments” (3 Kb chosen according to the average DNA fragment size with restriction enzyme HindIII digestion). Finally, based on the concept of chromosome territories ^6^, GEMs are then refined into purely intra-chromosomal “putative chromatin complexes”, ready for downstream analysis and visualization (Fig. 1b). ChIA-DropBox then enables visualization of the resulting chromatin interactions using several approaches. First, the data are converted to pairwise format and input files are generated to support existing tools: 2D contact map visualization using Juicebox ^7^ and loop visualization using BASIC Browser ^8^. In addition, ChIA-DropBox introduces a new web-based browser, named ChIA-View, for visualizing multiplex chromatin complexes. ChIA-View includes multiple display types, which support visualization of chromatin complexes in topological domains or around gene promoters (Fig. 1c).

### Quality Assessment of ChIA-Drop DNA sequencing data

First, ChIA-DropBox automatically generates a series of quality-assessment (QA) plots, every time the pipeline is run to automate assessment of each library (Fig. 2). The first set of QA plots are distributions of the insert sizes in the sequencing reads to assess the quality of library preparation, loading concentration, and sequencing. For example, these QA plots were generated for three libraries: a negative control library (empty droplets), a positive control library (pure-DNA) that fragments evenly distributing into droplets, and a ChIA-Drop library of chromatin complexes that distributing into droplets as a complex unit (Fig. 2a). Therefore, in the negative control, the droplets were predominantly empty. In the positive control, the droplets predominantly contained fragments for amplification. In the ChIA-Drop library, there were some empty droplets and some droplets that contained fragments for amplification. Thus, this QC plot indicates that the ChIA-Drop is a high-quality experimental library. While increasing the loading concentration could eliminate the empty droplets, it may also result in more droplets containing two independent chromatin complexes by random chance. Thus, a balance is necessary to obtain high-quality results, as indicated by the QA plot.

**Figure 2.**
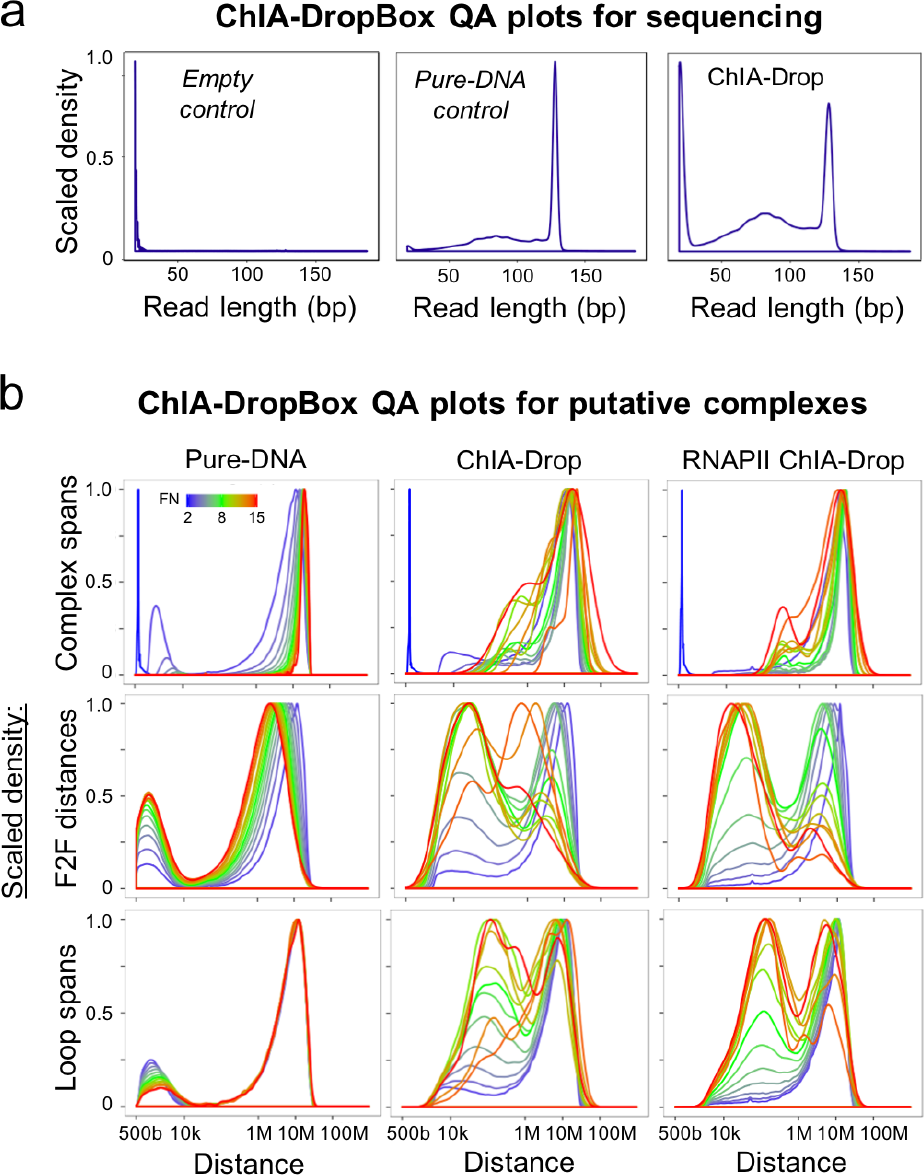
ChIA-DropBox automates quality assessment. **(a)** Quality-assessment (QA) plots for the raw sequencing library. For a negative control library of empty droplets (no chromatin material), the read length distribution has a peak at 20 bp, reflecting only the reads of DNA oligos and absence of chromatin DNA templates. For a positive control library of pure DNA fragments, the read length distribution has a peak around 130 bp, reflecting the full sequencing read-length (1×150 bp, minus library linker 20 bp) of pure DNA templates in all droplets. For a real ChIA-Drop library, the read length distribution has a peak at 20 bp (reflecting some empty droplets) and a peak around 130 bp (reflecting some droplets with chromatin templates). **(b)** QA plots of the distributions of various distance summaries of the putative chromatin complexes, as a function of the fragment number (FN) of the complex (color scale). The first row depicts the distribution of the spans of the chromatin complexes. The second row depicts the distribution of fragment-to-fragment (“F2F”) distances. The third row depicts the distribution of pairwise contact “loop” distances. Example QA plots are shown for three types of libraries. The first column is a control sample of pure-DNA fragments (no chromatin complexes). The second column is a ChIA-Drop sample. The third column is an RNAPII-enriched (immunoprecipitated) ChIA-Drop sample.

The second set of QA plots is used to assess the final chromatin complexes called from the ChIA-DropBox data processing workflow. These QA plots depict the distributions of various distance metrics used to summarize the complexes: (1) the span of the complex (from the first fragment to the last fragment), (2) the neighboring fragment-to-fragment (“F2F”) distances in the complex, and (3) the pairwise “loop” distances between all combinations of fragments in the complex (Supplementary Fig. 2). For example, ChIA-DropBox was used to generate these QA plots for three types of libraries in *Drosophila* S2 cells: a pure-DNA control sample (DNA was purified from the chromatin fragment after de-crosslinking), an unenriched ChIA-Drop sample, an RNAPII-enriched ChIA-Drop sample (Fig. 2b). In the pure-DNA control sample, most of the putative chromatin complexes have very long span (> 10 Mb in linear chromosomal distance) and long-range F2F distances (~ 1 Mb). Thus, the control sample of pure DNA reveals that the noise profile of ChIA-Drop is very long-range putative complexes, which likely resulted from independent DNA fragments being randomly partitioned into a shared droplet.

In contrast, the distance distributions for the ChIA-Drop sample and the RNAPII-enriched ChIA-Drop sample are much different than those for the control sample (Fig. 2b). For low-fragment-number chromatin complexes in the ChIA-Drop samples, most of the F2F distances are long-range (~1 Mb) indicating that these low-fragment-number GEMs are mostly noise, similar to the pure-DNA control sample. However, for high-fragment-number chromatin complexes in the ChIA-Drop samples, most of the F2F distances are in the biologically meaningful range of interaction distances (from 10 kb to 1 Mb), implying that these high-fragment-number chromatin complexes contain real signal for multiplex interactions. Thus, the QA plots from ChIA-DropBox indicate that the ChIA-Drop sample and the RNAPII-enriched ChIA-Drop sample are high-quality libraries containing real signal for multiplex chromatin interactions.

### Reconstruction and refinement of chromatin complexes

We next describe the detailed procedure for calling putative chromatin complexes (Fig. 3a). The initial steps of ChIA-DropBox data processing are: map reads to the reference genome, retain high-quality and uniquely mapped reads, parse the barcodes of the reads, and reconstruct the GEM droplets by grouping together reads with the same barcode. Once the GEM droplets are reconstructed, ChIA-DropBox refines them into putative chromatin complexes to attain interpretable data for analysis and visualization.

**Figure 3:**
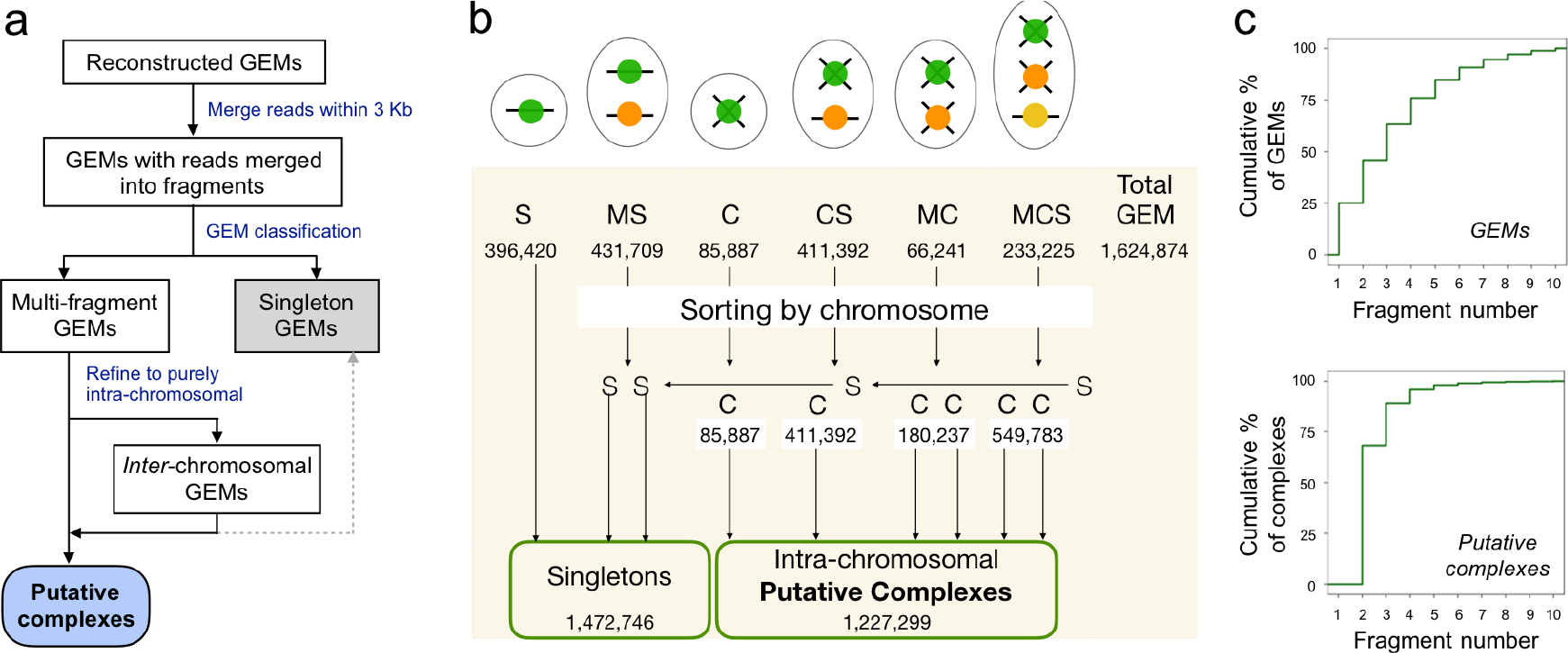
ChIA-DropBox reconstructs and refines putative chromatin complexes. **(a)** After attaining uniquely mapped reads and grouping together reads with the same barcode to reconstruct GEMs, ChIA-DropBox then has a comprehensive workflow for refining the GEMs into putative chromatin complexes. First, within each GEM, reads that overlap within 3 Kb in linear genomic alignment are merged into “chromatin fragments”. Second, GEMs are classified as multi-fragment GEMs or singleton GEMs. Based on the concept of chromosome territories, the multi-fragment GEMs are then refined to generate purely intra-chromosomal “putative chromatin complexes”. (**b**) Example breakdown of the results of applying the ChIA-DropBox workflow to a ChIA-Drop library. The schematic at the top illustrates the types of input GEMs for calling complexes: the colored circles indicate the chromosome of origin, and the number of fragments indicate singleton or multiple fragments (horizontal line or a cross). *S*: singleton; *MS*: multiple singletons; *C*: complex; *CS*: complex and singleton; *MC*: multiple complexes; *MCS*: multiple complexes and singleton(s). Below each category, the number of instances observed in the example library of *Drosophila* S2 cells is indicated. Also indicated are the total number of input GEMs, and the final number of putative complexes. (**c**) Cumulative distributions of the fragment number per GEM or per putative complex. There are many singleton GEMs and some high-fragment-number (inter-chromosomal) GEMs. In contrast, putative complexes predominantly contain between two and six fragments.

The first consideration is that during the experimental protocol, the linear amplification and barcoding initializes at random positions along each chromatin fragment. Thus, a single chromatin fragment could be represented by multiple reads with shifted positions and on different strands. Accordingly, within each computationally reconstructed GEM, ChIA-DropBox then assembles “chromatin fragments” by merging reads that have linear genomic alignments within 3Kb, consistent with the experimental chromatin fragment size.

Once reconstructed GEMs with merged chromatin fragments are ready, ChIA-DropBox then classifies and refines these GEMs into putative chromatin complexes to yield interpretable data (Fig. 3a). The goal of thus refinement step is to filter out two types of noise. The first type of noise is the presence of “singleton” chromatin fragments that are not in an interaction complex with any other fragments. A GEM may be comprised of only a singleton or of two or more singletons from different chromosomes (Fig. 3b). These purely singleton GEMs are filtered out. In addition, a GEM may contain a chromatin complex plus a singleton from a different chromosome (Fig. 3b). In this case, the GEM is refined by splitting it so that the singleton is filtered out and the putative chromatin complex is retained.

The second type of potential noise is a GEM that contains multiple chromatin complexes from different chromosomes (Fig. 3b). In this case, it is difficult to distinguish whether these inter-chromosomal multi-complex GEMs are true interactions or if they are examples of independent chromatin complexes from different chromosomes were randomly partitioned into the same droplet in the microfluidics device. Based on the concept of chromosome territories, the majority of chromatin complexes should be intra-chromosomal. Thus, ChIA-DropBox refines these GEMs by splitting them into purely chromosomal putative chromatin complexes (Fig. 3a). This refinement procedure was applied to a *Drosophila* ChIA-Drop library and the numbers of input GEMs by category are indicated, as well as the number of final putative chromatin complexes (Fig. 3b).

The results of calling putative chromatin complexes can be seen clearly by visualizing the distributions of the fragment number of the GEMs (before refinement) and the distribution of the fragment number of the putative chromatin complexes (Fig. 3c). Before refinement, around 25% of the GEMs are singletons, and there are also considerable GEMs with very high fragment number (reflecting inter-chromosomal multicomplex droplets). In contrast, after refinement, the putative chromatin complexes predominantly have between two and six fragments (Fig. 3c). Thus, ChIA-DropBox generates interpretable data from the raw ChIA-Drop FASTQ files by: mapping and filtering reads, reconstructing the GEMs based on the barcodes, and ultimately refining GEMs into putative chromatin complexes.

### Visualization for multiple chromatin interaction data

Once putative chromatin complexes are ready, ChIA-DropBox then automatically generates multiple output file formats to support diverse types of downstream analysis and visualization. First, the interactions from the putative chromatin complexes are converted to pairwise contacts (similar to Hi-C or ChIA-PET data), and the appropriate input files are generated to support existing analysis and visualization tools for pairwise contacts (Supplementary Fig. 3). Specifically, ChIA-DropBox calls Juicer ^9^ to generate a contact matrix for visualization of 2D contact maps using Juicebox ^7^ or Juicebox.js ^10^. In addition, since the ChIA-PIPE pipeline for ChIA-PET data processing has workflows for calling loops and calling binding peaks from pairwise contacts ^8^, ChIA-DropBox also generates a file format compatible with ChIA-PIPE data processing (see Methods). The resulting peaks and pairwise loops can then be visualized with high-resolution using BASIC Browser ^8^ (Supplementary Fig. 3).

While it is important to support existing pairwise visualization tools for comparison with ChIA-PET and Hi-C data, the fundamental advance of ChIA-Drop is that it captures multiplex chromatin complexes at the single-molecule level. Thus, ChIA-Drop data contain more detailed information about the true nature of chromatin complexes than can be displayed as pairwise contacts. Accordingly, ChIA-DropBox also introduces a new interactive browser, named ChIA-View, that is specifically designed to leverage the single-molecule and multiplex nature of ChIA-Drop data. ChIA-View is a fully featured and highly customizable browser that has four different display modes, each intended to facilitate its own type of downstream biological interpretation.

In ChIA-View, the fragments of each chromatin complex are displayed in their linear genomic alignments along the x-axis, and the different single-molecule complexes are arranged along the y-axis (Fig. 4). The first display mode is the “cluster view”. In the cluster view, the genomic region of interest is binned along the x-axis, and then the chromatin complexes with their binned fragments are arranged by similarity along the y-axis via hierarchical clustering (see Methods) (Fig. 4a). The cluster view enables biological interpretation by facilitating visualization of chromatin complexes across broad chromosomal regions, such as across multiple TADs (Fig. 4b). After hierarchical clustering has been applied to the binned chromatin complexes, the complexes can also be displayed with their original (unbinned) fragments (Fig. 4a). The display mode is named the “fragment view”. The fragment view is especially useful for visualizing the heterogeneity within topological domains at the single-molecule level (Fig. 4b). For example, fragment views of ChIA-Drop complexes in topological domains reveals substantial heterogeneity, consistent with recent reports from super-resolution microscopy ^11^.

**Figure 4:**
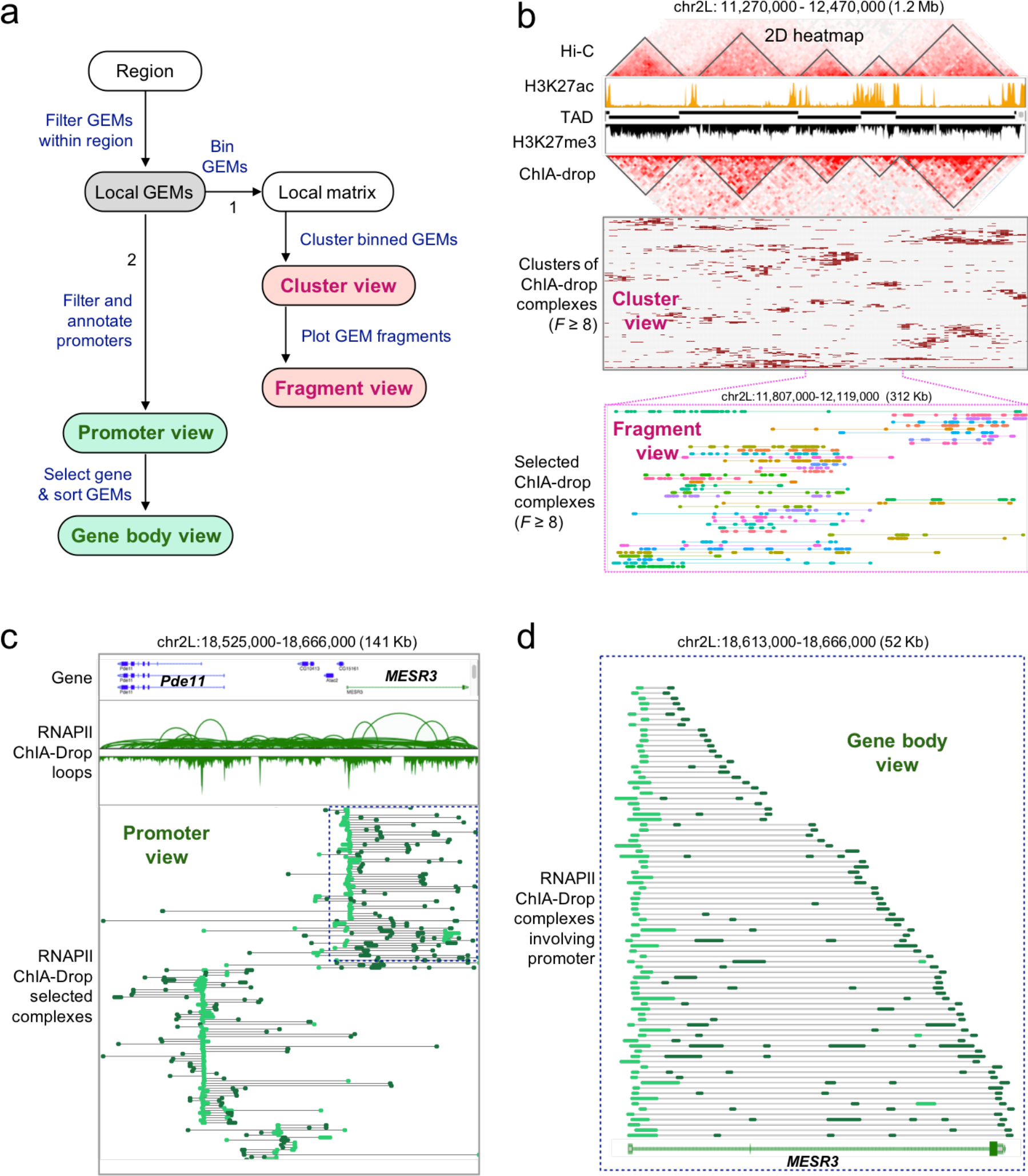
ChIA-DropBox introduces a new browser for visualizing multiplex single-molecule chromatin interactions. (**a**) A flowchart of ChIA-View. First, the user enters the coordinates of a genomic region of interest, and then GEMs mapping within this region are extracted. Once the “Local GEMs” have been extracted, there are two paths for visualization. The first path (“1”) is useful for visualizing unenriched ChIA-Drop data in topological domains, while the second path (“2”) is useful for visualizing RNAPII-enriched ChIA-Drop data around promoters. In path 1, the genomic region is binned, and then hierarchical clustering is applied to arrange the GEMs by similarity along the y-axis (“Cluster view”). Once the GEMs have been arranged, the original un-binned data can be viewed with the same y-axis arrangement (“Fragment view”). In path 2, GEMs involving promoters are extracted and the fragments in each GEM that overlap promoters are highlighted in a bright color (“Promoter view”). Then, an individual gene within the region can be selected, and GEMs involving that gene’s promoter are sorted by GEM span (“Gene body view”). (**b**) An example of multiple visualization types for unenriched ChIA-Drop data in a 1.2 Mb region in *Drosophila* S2 cells. First, Hi-C data, ChIP-seq data, and domain calls (TAD) are displayed as a reference. Then, a 2D heat map view of ChIA-Drop data is shown. Below that, a ChIA-View “Cluster view” of the same region is shown. Below that, a magnified view of a single TAD is shown with ChIA-View “Fragment view”. (**c**) An example of the ChIA-View “Promoter view” of RNAPII-enriched ChIA-Drop in a 141 Kb window containing the gene *Pde11* and *MESR3*. Gene annotations, RNA-seq data, and pairwise ChIA-Drop loops are shown for reference. (**d**) The gene *MESR3* is selected, and a zoomed-in ChIA-View “Gene body view” is displayed.

While the cluster view and fragment view in ChIA-View are well suited for unenriched ChIA-Drop data, the next two display modes are specifically designed for RNAPII-enriched ChIA-Drop data and take into account the promoter-centric nature of the data. In the “promoter view”, chromatin complexes within the genomic region are filtered to retain those that overlap promoters (Fig. 4a). Then, the complexes are grouped together along the y-axis based on which gene’s promoter they overlap. Within each complex, the fragments are color coded to indicate whether they overlap a promoter element or a non-promoter element. For example, promoter views of RNAPII-enriched ChIA-Drop data in *Drosophila* reveals that most single-molecule complexes involve only one promoter and multiple non-promoter elements (Fig. 4c). This observation is consistent with a recent report from super-resolution microscopy that 80% of RNAPII foci in the nucleus have a single RNAPII protein ^12^. Once the promoter-involving complexes have been filtered and color annotated, complexes within a single gene body can then be viewed. In the “gene body view” display mode, chromatin complexes fully contained within the body of a single gene are retained and then they are sorted by span along the y-axis (Fig. 4a). For example, the gene body view of RNAPII-enriched ChIA-Drop data in *Drosophila* reveals a pattern that would be consistent with a one-side extrusion model of transcription (Fig. 4d). ChIA-View is available as an easy-to-use *R* Shiny application (Supplementary Fig. 4).

## Discussion

In conclusion, ChIA-DropBox is a major advance both in terms of the processing and the visualization of a novel data type. Considering that ChIA-Drop experiments use pooled sequencing of barcoded fragments, the computational reconstruction of the data is essential to yield interpretable results. Thus, ChIA-DropBox is a crucial companion of ChIA-Drop. Starting with raw sequencing reads, it performs read mapping, barcode parsing, GEM reconstruction, and refinement of putative chromatin complexes.

For visualization, ChIA-DropBox automatically converts the data to pairwise format and generates input files to support Juicebox views of 2D contact maps and BASIC Browser views of loops. ChIA-DropBox also introduces a new browser named ChIA-View, the first visualization tool for single-molecule multiplex chromatin complexes to the best of our knowledge.

Overall, the current version of ChIA-DropBox including ChIA-View cover all the core elements necessary for processing, analyzing, and visualizing the multiplex chromatin interaction data from ChIA-Drop experiments. Notably, ChIA-View was designed to support visualization of any multiplex chromatin interaction data, and can readily support the visualization of multiplex interactions from GAM or SPRITE. Thus, we envision ChIA-View as a comprehensive browser for multiplex interactions, which will also enable comparative analyses of the data generated from different experimental methods. In addition, future versions of ChIA-DropBox will be extended to support processing of data generated by GAM and SPRITE.

The introduction of ChIA-DropBox will enable researchers to immediately begin interpreting their ChIA-Drop experiments. As there are many outstanding questions in the field of 3D genome organization that can only be addressed at the single-molecule level, we anticipate that these computational advances and tools will have high visibility and high impact. Empowered with ChIA-DropBox and ChIA-View, researchers will be able to explore many crucial topics, such as: the cell-to-cell variation of chromatin interactions in topological domains; the mechanisms of chromatin loop and domain formation; the landscape of chromatin interactions in heterogeneous cell populations, such as tumors’ and immune cell populations; and the multiplex nature of transcriptional regulation and gene-gene interactions.

## Acknowledgements

This study is supported by a Jackson Laboratory Director’s Innovation Fund (DIF19000-18-02). Y.R. is funded by 4DN (U54 DK107967) and ENCODE (UM1 HG009409) consortia. Y.R. is also funded by Human Frontier Science Program (RGP0039/2017), and supported by Florine Roux Endowment.

## Author contributions

S.Z.T developed the ChIA-DropBox pipeline and D.C., M.K. B.L., M.Z., and Y.R. provided input. S.Z.T. developed ChIA-View and D.C., M.Z., and Y.R. provided input. D.C. developed the utility to make the output compatible with ChIA-PIPE loop calling. S.Z.T and D.C. created the figures and M.Z. and Y.R. provided input. D.C., S.Z.T., and Y.R. wrote the manuscript. All authors read and approved the final manuscript.

## Competing interests

The authors do not declare any competing interests.

## Methods

### ChIA-DropBox workflows

ChIA-DropBox includes two components: the data-processing and the data-visualization. The data-processing workflow contains four categories of procedures: read alignment; GEMcode identification and read filtering; GEM calling; and chromatin-complex calling. There are 39 processing steps distributed among these four categories. At the end of GEM calling, ChIA-DropBox has a step to generate multiple formats of files for various purposes, including quality-assessment (QA) plots, and statistics tables. In the data visualization workflow, three types of visualization are supported. First, ChIA-DropBox converts the multiplex chromatin interaction data to pairwise format and automatically generates input files to visualize 2D contact maps using Juicebox ^7,9^. Second, ChIA-DropBox uses the pairwise-format data to generate input files for presenting interaction loops in genome browsers. Third, ChIA-DropBox includes a novel web-based browser, named ChIA-View, for visualizing multiplex chromatin interactions.

### Read alignment

In this step, reads are aligned to the reference genome to generate a BAM file of aligned reads with GEMcodes. In addition, a statistics table is generated, which includes library name, total paired-end read count, PCR redundancy rate, among others.

First, base-calling is performed using the 10X Genomics *longrange mkfastq* pipeline (v2.1.5), and then a reference genome index for the *longranger* pipeline is created using *longrange mkref* (v2.1.5). Next, mapping is performed via the 10X Genomics *longrange wgs* pipeline (v2.1.5) (see Supplementary Methods). When using *longranger wgs*, the parameters (e.g., number of processors; amount of memory) are set in the “MRO” file (see Supplementary Methods). Important values in the MRO file are the library ID, the path and names of the FASTQ files, and the path of the reference genome folder (see Supplementary Methods).

### GEMcode identification and read-quality filtering

In this step, the GEMcode of each read is identified, the read ID is renamed to include the GEMcode, and the reads are filtered for quality.

First, GEMcodes are extracted from the BAM file by accessing the “BX” tag field in the BAM files. Then, the ID of each read is renamed to include the GEMcode, along with its numeric GEM ID (see Supplementary Methods). This re-naming allows the GEM ID to be propagated forward when the BAM files are converted to other output formats for downstream analyses.

Second, reads are filtered for quality. The R1 read of each pair is retained for downstream analysis, since it contains both the GEMcode and the amplified DNA sequence. After removing non-primary alignments and low map-quality reads (MAPQ < 30 and read length < 50bp), each read is then extended by 500 bp from its 3’ end, consistent with the DNA template length in the ChIA-Drop experiment. These filtering steps are performed using *pysam* (v0.7.5) in *python* (v2.7.9) and *samtools* (v0.1.19).

### Calling GEMs

In this step, GEMs are reconstructed. Uniquely mapped reads with the same GEMcode are grouped together. Overlapping reads within each GEM are merged into chromatin fragments. Then, the GEMs are refined and classified.

All reads with the same GEMcode and their extended genomic coordinates overlapping within 3Kb distance are merged to represent a chromatin fragment, using the *python* package *pybedtools* (v0.7.10) ^13^. This distance was chosen because the average fragment length of the genomic DNA in *Drosophila* after HindIII digestion is 3 Kb. This merging step also collapses any duplicate reads, so it is not necessary to include a separate PCR de-duplication step. Three output file formats of the GEMs for the downstream analysis are 1) one GEM per line, 2) one fragment per line, and 3) one read per line.

After merging overlapping reads into chromatin fragments, each GEM is then categorized based on the number of fragments it contains. A GEM with one chromatin fragment (*F* = 1) is considered a singleton GEM, whereas a GEM with two or more fragments (*F* ≥ 2) is considered a multiplex GEM, representing a chromatin interaction complex with multiple fragments. Based on the genomic coordinates of the chromatin fragments, multiplex GEMs are further classified as intra-chromosomal GEMs if all fragments within a GEM mapped to the same chromosome, or inter-chromosomal GEMs if the fragments within a GEM mapped to different chromosomes. Since most interactions occur within chromosome territories ^6^, the inter-chromosomal GEMs are then split into their intra-chromosomal sub-components. After the refinement and classification, an additional output file is generated with one “sub-GEM” per line.

### Statistic report table

Extensive quality-assessment statistics are generated, such as the total read count, the PCR redundancy rate, and the number of mappable reads with GEMcodes. The read count is the reported after each filtering step: number of uniquely mapped R1 reads, reads with mapping score ≥ 30, and reads with length <50 bp.

Statistics are also reported for the GEM calling and classification steps. The values reported include: the original GEM number; the number of GEMs with a single fragment; and the number of GEMs with multiple fragments. The number of multi-fragment GEMs is then further dissected into the number of intra-chromosomal GEMs and inter-chromosomal GEMs, as well as the number of sub-GEMs after splitting the inter-chromosomal GEMs.

### Quality-assessment plots

ChIA-DropBox also generates quality-assessment plots to help understand the data. In addition to the aforementioned QA plots, plots are generated to depict distributions of read length, fragment lengths, read count per fragment, neighboring-fragment distances, and the fragment number per GEM (see Supplementary Methods).

### Data visualization

To visualize multiplex chromatin interaction complexes in a genome browser-based view, we developed a linear multi-fragment view tool, called ChIA-View, which displays the chromatin fragments in either a clustering heatmap view, or a multi-fragment linear alignment view. In a standard ChIA-View, along the y-axis, GEMs are arranged by similarity, which is computed via hierarchical clustering (using the *R* package *pheatmap* (v1.0.10) with “clustering_distance_rows” = “euclidean” and “clustering_method” = “complete” in default). ChIA-View also includes variant modes: PE ChIA-View for annotating promoter and non-promoter fragments, and Gene body ChIA-View for sorting GEMs to demonstrate RNAPII extrusion model.

### ChIA-View for unenriched data: “Cluster” mode and “Fragment” mode

The input file for a standard ChIA-View is a REGION file, which is a per-line-per-fragment data from the ChIA-DropBox processing pipeline. The ChIA-View workflow is performed as follows. First, all fragments within a given genomic coordinate are selected by *bedtools* (v2.27.1). Second, GEMcodes specific fragments are assigned to 100 bins per GEM in default (or a fix length bin), constructed a Boolean matrix (per row per GEM, and per column per bin). Continuously, the Boolean matrix’s GEM order (row order) was determined by hierarchical clustering, and plot out by *R* package *pheatmap* (v1.0.10) in clustering heatmap views. Finally, the original GEMs were plotted according to alignment order achieved from the hierarchical clustering function. In the resulting visualization, the entire GEM span is indicated with a thin line, and the individual fragments are displayed with thick lines (linear alignment views of ChIA-View). Each row is a GEM and the GEMs are displayed in assorted colors. The linear alignment view of ChIA-View is performed using *R* (v3.5.1) with mainly *ggplot2* (v2_3.0.0).

Other *R* packages used in ChIA-View include: *pheatmap* (v1.0.10), *ggplot2* (v3.0.0), *dplyr* (v0.7.6), *bedr* (v1.0.4), *ggforce* (v0.1.3), *cowplot* (v0.9.3), *gtable* (v0.2.0), *gridExtra* (v2.3), *reshape2* (v1.4.3), *RColorBrewer* (v1.1-2), *ggrepel* (v0.8.0), and *ggpubr* (v0.1.7).

### ChIA-View for enriched data: “Promoter” mode and “Gene body” mode

For a RNAPII ChIA-Drop data, we are interested in interplay among regulatory elements such as promoters and enhancers. Thus, we annotate each fragment as a promoter fragment if its coordinates overlap with a gene promoter region (TSS ± 250 bp); all other fragments are denoted as non-promoters. Fragments are color-coded by this classification.

The Gene body ChIA-View starts from a promoter annotated REGION file used for Promoter ChIA-View, and zooms into a particular gene. Then sorting GEMs order by a self-defined method and plot out, instead of hierarchical clustering in standard ChIA-View. This sorting method selects GEMs that owning fragments within the promoter region of the selected gene, then sorting Gems by their local GEM coverage from short to long.

### Code availability

The ChIA-DropBox data processing pipeline is publicly available at: https://github.com/TheJacksonLaboratory/ChIA-DropBox.git. The ChIA-view visualization tool for multiplex interactions is publicly available at: https://github.com/TheJacksonLaboratory/ChIA-view.git. Both repositories contain README files with further details on how to get started using the software.

### Data availability

The ChIA-Drop and RNAPII ChIA-Drop data showcased in this paper were publicly released previously (raw sequencing FASTQ files; processed data files of aligned reads with parsed barcodes; and final files of chromatin interaction complexes) via the Gene Expression Omnibus under SuperSeries accession number: GSE109355.

---

**Supplementary Figure 1:**
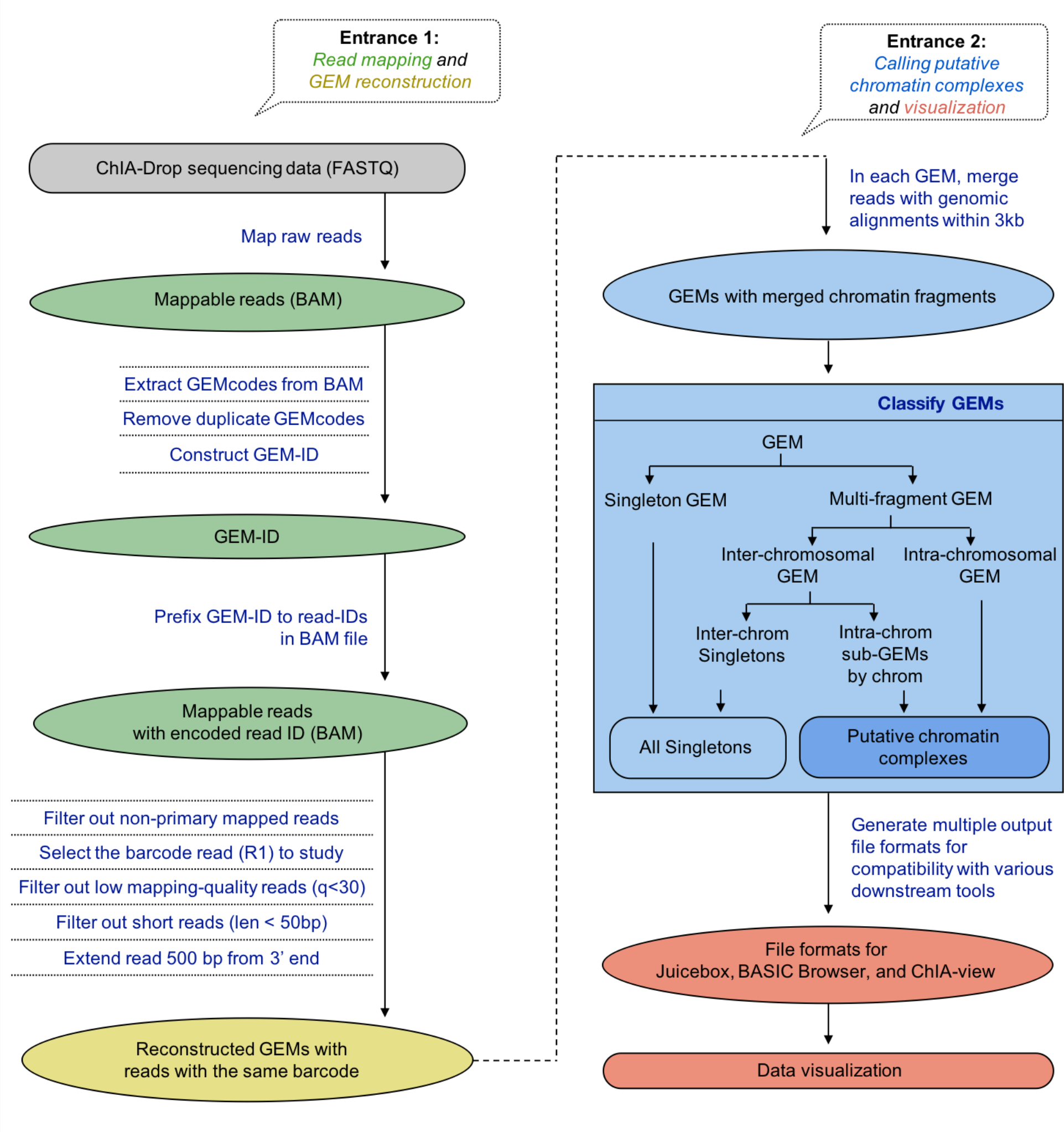
Detailed flowchart of ChIA-DropBox data processing. A flowchart of the detailed steps in the ChIA-DropBox processing pipeline. Gray oval shapes represent data files. *Green*: FASTQ files of sequencing reads are mapped to the reference genome to generate a BAM file. The barcode is extracted from each sequencing read and then prefixed to the read ID in the BAM file. Reads are filtered to retain high quality reads; the R1 read, which contains both the barcode and genomic sequence, is retained; and the read is extended. *Yellow*: GEMs are reconstructed by grouping together reads with the same barcode. *Blue*: GEMs are further refined. First, within each GEM, reads that have genomic alignments within 3 Kb of each other are merged into “chromatin fragments”. A refinement procedure is applied to obtain a set of multiplex and intra-chromosomal “putative chromatin complexes”. *Red*: the putative chromatin complexes are output in multiple file formats to enable downstream analysis and visualization. First, the data are converted to pairwise format and input files are generated for viewing 2D heat maps in Juicebox and viewing loops in BASIC Browser. Then, an input file is generated for viewing multiplex interactions in ChIA-View.

**Supplementary Figure 2.**
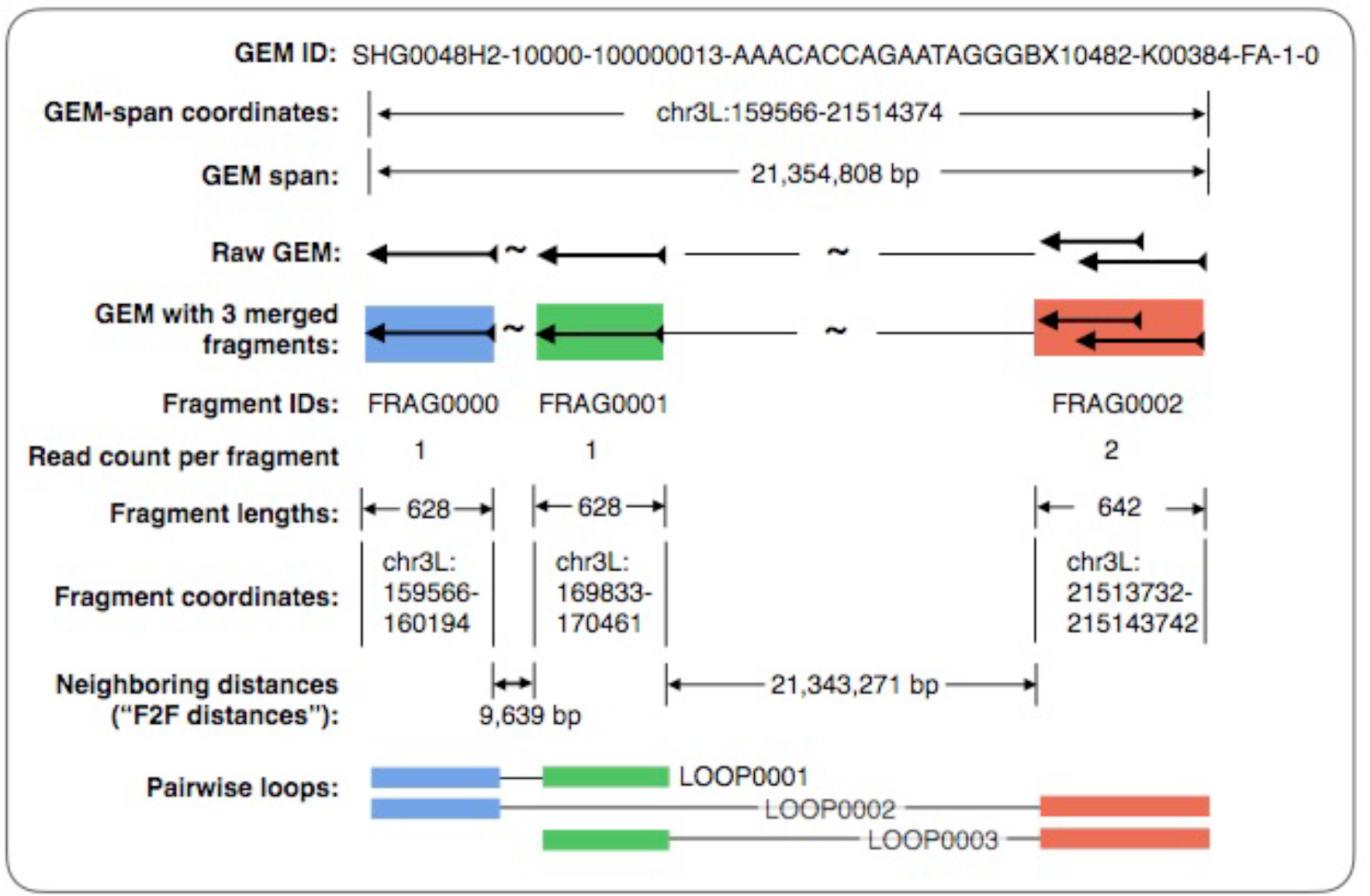
ChIA-DropBox data formats and terminology. Each intra-chromosomal GEM is given a unique identifier (ID), which also encodes information about the GEM (e.g., the sequencing run, and the number of fragments in the GEM; see Methods). Each GEM has an outer span from its leftmost fragment to its rightmost fragment, and corresponding coordinates of the GEM-span. After a length extension, reads in the same GEM located within 3Kb of each other are merged into “chromatin fragments”. Each fragment has an ID, a read number, a length, and coordinates. The distances between fragments in a GEM are considered in two ways. First, the distances between neighboring fragments are considered (fragment-to-fragment, or “F2F”, distances). Second, the GEM is converted to pairwise loops and the pairwise distances are considered.

**Supplementary Figure 3:**
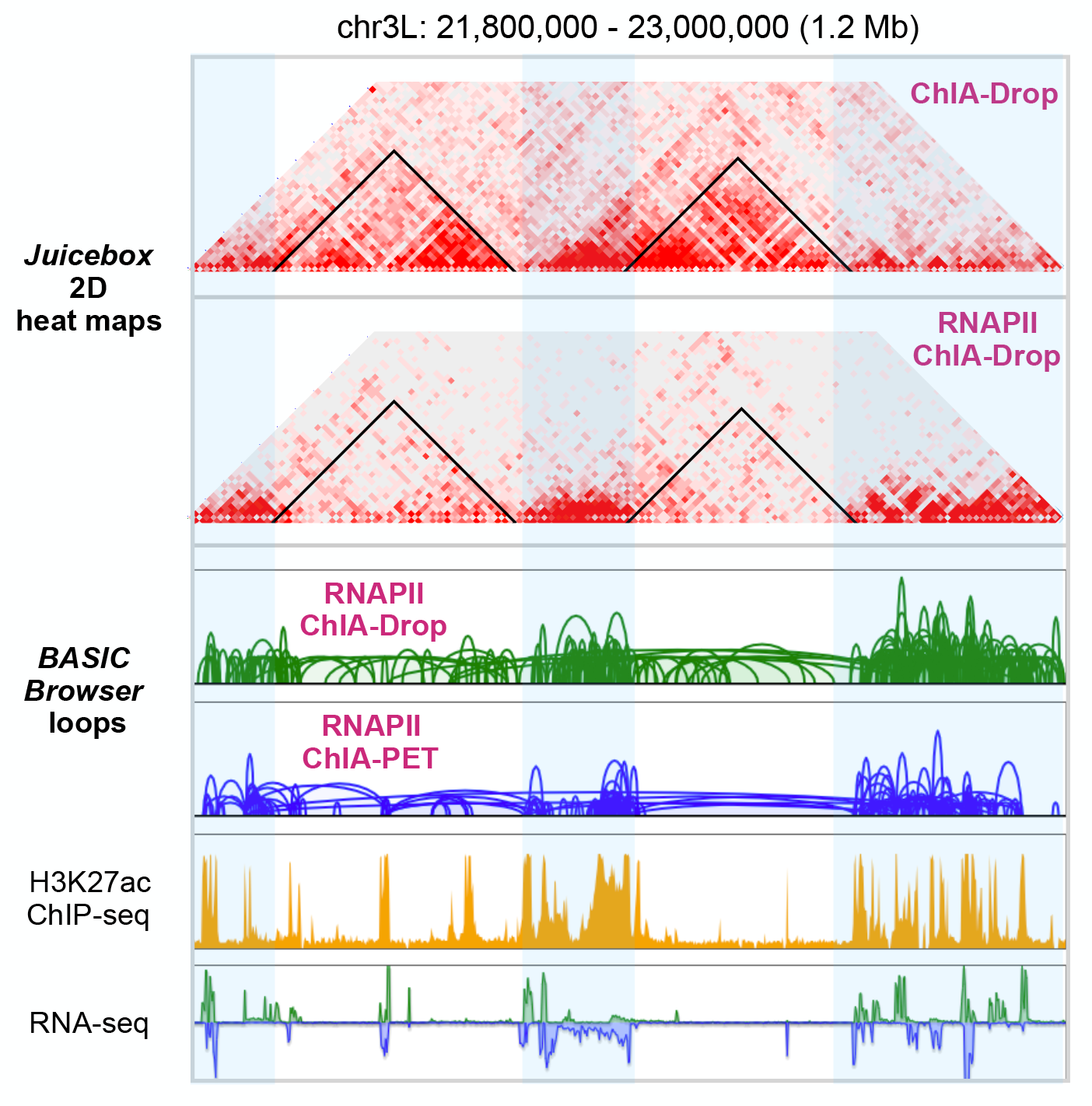
ChIA-DropBox automatically converts the putative chromatin complexes to pairwise format and generates input files for existing visualization tools. After ChIA-DropBox reconstructs and refines the putative chromatin complexes, it then supports existing visualization tools by automatically converting the data to pairwise format and generating the appropriate file formats. A contact matrix is generated for viewing 2D heatmaps in Juicebox. An input file is generated for performing loop calling via ChIA-PIPE and then viewing the loops in BASIC Browser. Juicebox 2D heat maps of an unenriched ChIA-Drop sample and an RNAPII-enriched ChIA-Drop sample in *Drosophila* S2 cells. In *Drosophila*, ChIA-Drop signal is enriched in the transcriptionally inactive topological domains, while RNAPII-enriched ChIA-Drop signal is enriched in the transcriptionally active gap regions between domains. BASIC Browser views of RNAPII-enriched ChIA-Drop loops and RNAPII-enriched ChIA-PET loops (both sets of loops called using ChIA-PIPE). BASIC Browser also displays a ChIP-seq coverage track for an active histone mark and an RNA-seq track. Loops are enriched in the gap regions between domains for both RNAPII-enriched ChIA-Drop and RNAPII-enriched ChIA-PET.

**Supplementary Figure 4:**
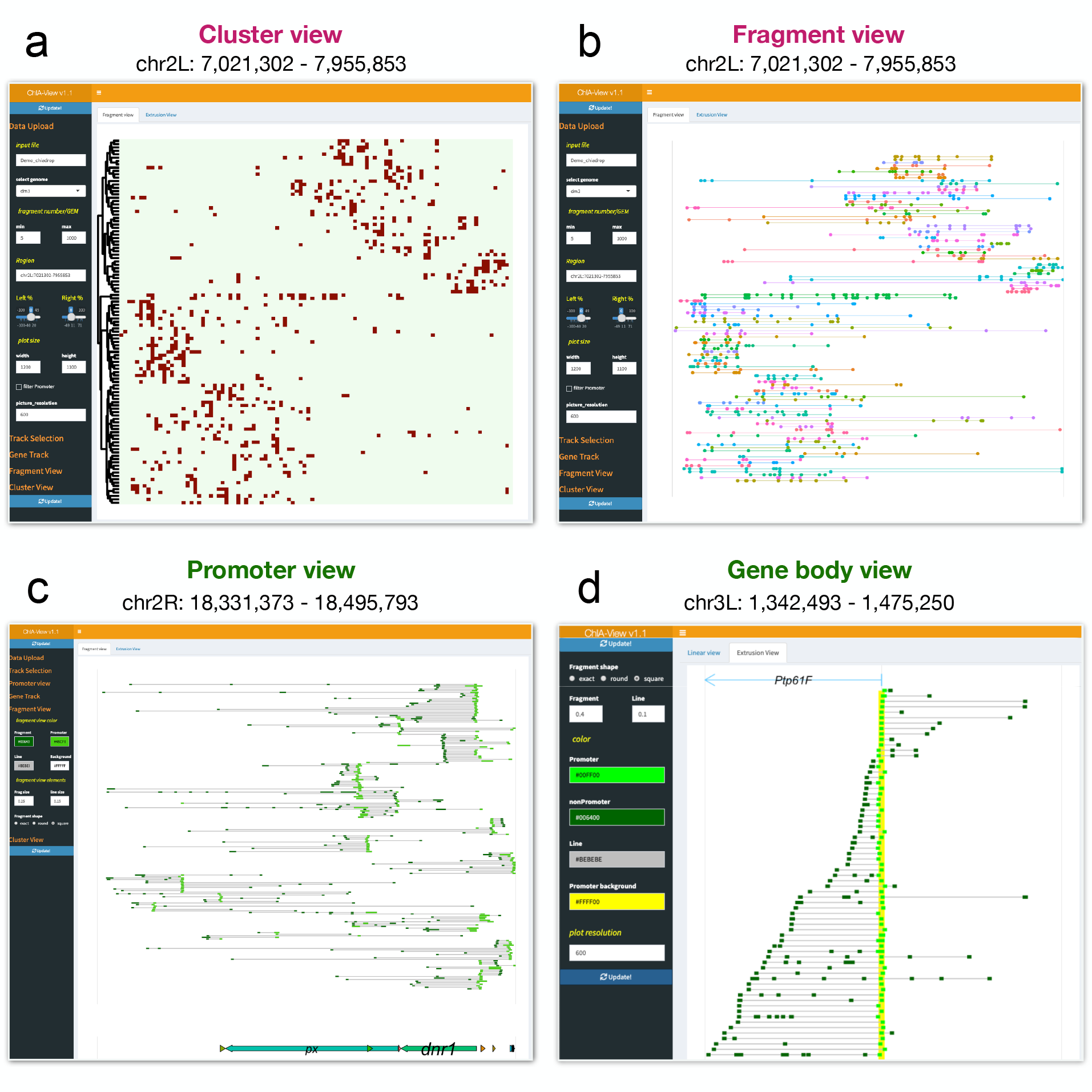
ChIA-View is a new web-based *R* Shiny app for visualizing multiplex chromatin interactions. ChIA-View is an easy-to-use *R* shiny app for high-resolution visualization of single-molecule multiplex chromatin interactions. ChIA-View facilitates biological interpretation through four different display modes: (a) “Cluster view”, (b) “Fragment view”, (c) “Promoter view”, and (d) “Gene body view”. In the ChIA-View browser, there is core control panel on the left side for navigation. Using this control panel, the user can upload data (or select demo data pre-loaded in the browser). Once a data set is selected, the user can then: select a display mode; enter the coordinates of a genomic region of interest; filter GEMs by fragment number; refine properties of the display (e.g., colors, line width, dimensions); and export the display to a high-resolution, publication-quality image file. The “Cluster view” and the “Fragment view” are especially well suited for unenriched ChIA-Drop data. The “Cluster view” displays binned chromatin complexes over broad chromosomal regions, and the different complexes are arranged by similarity along the y-axis via hierarchical clustering. The “Fragment view” retains the y-axis arrangement from the “Cluster view”, but instead displays the original chromatin fragments. The “Promoter view” and the “Gene body view” are particularly well suited for RNAPII-enriched ChIA-Drop data. The “Promoter view” filters and retains chromatin complexes overlapping promoters, and then highlights the promoter-involving fragments in a different color. The “Gene body” view displays the chromatin complexes involving the promoter of a specified individual gene, which the chromatin complexes arranged by span.

## Supplementary Methods

The ChIA-DropBox pipeline includes two parts: data processing and data visualization. The data-processing workflow includes four main categories: (1) read mapping and barcode identification; (2) GEM reconstruction; (3) refinement of GEMs into putative chromatin complexes; (4) generating various file formats to support downstream analysis and visualization; and (5) quality assessment. The data-visualization workflow includes support for existing visualization tools and introducing a new visualization tool. For existing visualization tools for pairwise interaction data, ChIA-Dropbox generates a contact matrix for viewing 2D heat maps in Juicebox and generates a file for performing loop calling via ChIA-PIPE, with subsequent visualization of the loops in BASIC Browser. ChIA-DropBox also introduces a new browser, named ChIA-View, specifically designed for visualizing single-molecule multiplex chromatin interactions.

### Read mapping

After the droplet-barcoded and pooled ChIA-Drop amplicons are sequenced on a standard Illumina platform (e.g. Miseq, NextSeq 500, or HiSeq 4000), ChIA-DropBox begins data processing by performing read mapping via the 10X Genomics Chromium software suite.

First, the 10X Genomics ‘longranger mkfastq’ (v2.1.4) command is used with default parameters to convert the data set of barcoded reads to demultiplexed FASTQ files. Then, 10X Genomics ‘longranger wgs’ (v2.1.5) is used in whole-genome mode to map reads to the reference genome, with additional parameters specified as suitable for the computing environment (e.g. “longranger wgs $LIB MCP.mro --jobmode=local --localcores=32 --localmem=150gb”). It is important to set the correct information in the ‘MCP.mro’ file, and these can be achieved using a utility script in ChIA-DropBox named ‘MCP.init.sh’. Specifically: (I) ‘sample_id’ is library name; (II) “gem_group” is the library specific digits code, which will appear at the end of ‘BX’ tag of BAM files (e.g., in a ‘BX’ tag ‘BX:Z:AAACCCAAGAAGTGAG-10482’, the ‘gem_group’ is ‘10482’); (III) “read_path” is the absolute path to the FASTQ directory; (IV) “sample_names” is the prefix of FASTQ file names (e.g., for FASTQ file named “SHG0048H2_GT18-08819_SI-GA-A2_S30_L007_R1_001.fastq.gz”, the “sample_names” is “SHG0048H2_GT18-08819_SI-GA-A2”); (V) ‘sex’ should be set to ‘m’ (male) or ‘f’(female) according to the cell line; and (VI) ‘reference_path’ is the absolute path to the reference genome file, which could be downloaded from 10X Genomics website, or generated using the ‘longranger mkref’ command.

The runtime for this step is typically 3-8 hours (e.g. for a lane of Hiseq sequencing data), after which a BAM file of mappable reads with barcode information is generated.

### Barcode identification

First, barcodes are extracted from the BAM file by accessing the “BX” tag field in the BAM files using the *pysam* module (v0.7.5) in *python* (v2.7.9). For simplicity, each unique barcode sequence is assigned an identifier (ID) as follows. First, the list of barcodes is deduplicated to obtain a list of unique barcodes. Then the unique barcodes are sorted alphabetically and are assigned an incrementing numerical ID starting from “100,000,000”. Then, an ID is generated for each GEM by concatenating the library ID, the barcode numeric ID, and the barcode sequence (each separated by a dash symbol (“-”)). An example of a GEM-ID is “SHG0048H2-10000-100000944-AAACCCAAGAAGTGAGBX10482”, in which “SHG0048H2” is the library ID, “10000” is a sub-group ID, “100000944” is the barcode numeric ID, “AAACCCAAGAAGTGAG” is the barcode sequence, “BX” is the BAM tag containing the barcode, and “10482” is the “gem_group” (library numeric ID).

Based on these GEM IDs, the read IDs of mappable reads in the BAM files are then renamed using the *pysam* module (v0.7.5) in *python* (v2.7.9). This re-naming allows the GEM-ID to be propagated forward when the BAM files are converted to BED files for downstream analysis. Specifically, the read ID is modified by prefixing it with the relevant GEM ID. For example, a new read ID could be: “SHG0048H2-10000-100000944-AAACCCAAGAAGTGAGBX10482-K00384:133:HTYWJBBXX:7:2224:26707:29729”, which includes the GEM ID (“SHG0048H2-10000-100000944-AAACCCAAGAAGTGAGBX10482”), and the original read ID (“K00384:133:HTYWJBBXX:7:2224:26707:29729”).

### Quality filtering of the data

Next, ChIA-DropBox performs a quality-filtering step to retain only the high-quality data. Reads classified as “not primary alignment” or “supplementary alignment” are discarded by using the “-F 2304” flag in *samtools view* (v0.1.19). For convenience in subsequent steps, the data are then converted from binary format (BAM file) to text format (paired BED files) using *bedtools bamtobed* (v2.27.0). For simplicity, R1 reads are retained for subsequent steps, as they contain the barcode and the genomic sequence (R2 reads do not contain the barcode). R1 reads are further filtered by requiring MAPQ ≥ 30, and very short reads from multiple displacement amplification (MDA) bias are filtered out by requiring length ≥ 50 bp. After this filtering, a mean read length of 120-130 bp is typically obtained. Next, each read is extended by 500 bp from its 3’ end, reflecting the DNA template size that was used for sequencing (~ 300 - 600 bp).

### Reconstructing GEMs and refining putative chromatin complexes

The goal of this step is to reconstruct the chromatin droplets and then refine them into putative chromatin complexes. First, chromatin droplets are reconstructed by grouping together reads with the same barcode (and thus the same GEM ID). The resulting intermediate file format referred to as a “BCline” file has one GEM per line with further information about which reads the GEM contained.

Next, the individual reads within each GEM are merged into chromatin fragments. Since chromatin fragments are randomly primed for linear amplification (ChIA-Drop experimental protocol), the same chromatin fragment could be represented in the ChIA-Drop data by multiple different but overlapping sequencing reads. Therefore, all reads with same GEM ID whose extended genomic coordinates overlap within 3 kb distance are merged to represent a chromatin fragment (using the *python* package *pybedtools* (v0.7.10)). This distance was chosen because the average fragment length of the genomic DNA after cutting with HindIII is 3-4 kb. This merging of reads into chromatin fragments also de-duplicates PCR redundancies *de facto*, so it is not necessary to perform a separate de-duplication step.

Once GEMs with merged chromatin fragments are obtained, three types of output files are generated: 1) “GEMLINE” (one GEM per row with information about the fragments in each GEM), 2) “FRAGLINE” (one chromatin fragment per row with information about the reads in each chromatin fragment), and 3) “READLINE” (one read per line with a pointer to its chromatin fragment). These 3 files can be seen as the first layer outputs of ChIA-DropBox, and are used for generating quality-assessment statistics and refining putative chromatin complexes. They are also useful to trace back the source reads belong to a chromatin fragment in a GEM.

After merging overlapping reads into chromatin fragments, GEMs are refined into putative chromatin complexes in multiple steps. First, individual GEMs are categorized based on the number of fragments they contain. A GEM with one chromatin fragment (*F* = 1) is considered a singleton GEM, whereas a GEM with two or more fragments (*F* ≥ 2) is considered a multiplex GEM, representing a chromatin interaction complex with multiple fragments. Next, based on the genomic coordinates of the chromatin fragments, multiplex GEMs are further classified as intra-chromosomal GEMs if all fragments within a GEM are mapped to the same chromosome, or inter-chromosomal GEMs if the fragments within a GEM mapped to different chromosomes.

Considering the concept of chromosome territories, the goal is then to obtain a set of high-confidence intra-chromosomal multiplex chromatin complexes, as the inter-chromosomal GEMs could potentially be instances where independent chromatin complexes were randomly partitioned into the same droplet. Thus, inter-chromosomal GEMs are split into purely intra-chromosomal sub-GEMs. For instance, a given GEM possesses 6 chromatin fragments, in which 2 were mapped to one chromosome and the other 3 were to another chromosome, 1 was mapped to a third chromosome. This inter-chromosomal GEM will be split into two intra-chromosomal subsets and one single-fragment subset (singleton). Thus, this procedure refines the GEMs with merged chromatin fragments into putative chromatin complexes to be used in downstream analysis and visualization.

### Output in diverse file formats

Once putative chromatin complexes have been refined, ChIA-DropBox generates output files in multiple formats to accommodate a diverse set of downstream analysis and visualization. First, the file of putative chromatin complexes is converted to GFF format, which allows linked fragments in each chromatin complex to be visualized in IGV as a genome ideogram track. Second, multiple types of coverage tracks (bedgraph) files are generated to enable assessment of the binding and enrichment and coverage. Coverage tracks are generated separately based on the read pileups, the fragment pileups, and the chromatin complex pileups.

To support existing visualization tools for pairwise chromatin contacts, ChIA-DropBox automatically converts the putative chromatin complexes to a file of pairwise contacts (BEDPE). This BEDPE file can then be used as input for generating a 2D contact map to view in Juicebox and as input for calling loops with ChIA-PIPE to view in BASIC Browser (described in more detail in a later section). In addition, ChIA-DropBox generates a “REGION” file, which is the source file for loading data into the new ChIA-View multiplex browser introduced here (described in detail in a later section).

### Quality Assessment statistics

Next, key summary statistics are gathered from the output of each step of the pipeline and a quality-assessment statistics table is generated. Based on the mapping and barcode identification steps, the following summary statistics are included in the QA table: the total read count; the PCR redundancy rate; the number of reads that are uniquely mapped and have a barcode; the number of reads that pass quality filtering (MAPQ ≥ 30 reads and length ≥ 50 bp); and the number of unique barcodes. Based on the GEM reconstruction and putative chromatin complex refinement steps, the following summary statistics are also recorded: the total GEM number; the number of GEMs with a single chromatin fragment (singleton); the number of GEMs with multiple chromatin fragments; the number of intra-chromosomal multiplex GEMs; the number of inter-chromosomal multiplex GEMs; and the final number of putative chromatin complexes after refinement.

### Visualization of ChIA-Drop data in pairwise format

As described above, once the putative chromatin complexes have been obtained, ChIA-DropBox automatically converts the data to pairwise format (BEDPE file). To support 2D heatmap visualization, ChIA-DropBox then uses Juicer to convert the file of pairwise interactions into a contact matrix file (.hic file). The .hic file can then be visualized directly in Juicebox or Juicebox.js as a 2D contact map.

In addition, the file of pairwise interactions is converted to a format that allows loop calling to be performed using ChIA-PIPE. Specifically, the BEDPE file and reference genome FASTA file are used to create “pseudo paired-end FASTQ files” by extracting the sequences of interacting fragments from the reference genome. These pseudo paired-end FASTQ files are then processed by ChIA-PIPE, such that the loops visualized for ChIA-Drop data are directly comparable with the loops visualized for ChIA-PET data.

Briefly, ChIA-PIPE calls loops as follows. First, paired-end tags (PETs) are aligned to the reference genome and only uniquely-mapped, non-redundant PETs are retained. Then, each tag is extended by 500 bp in its 5’ direction and PETs that have both ends (R1 and R2) overlapping are merged into loops. The interaction frequency of each loop is then the number of PETs that contributed, and the loops can then be visualized in BASIC Browser and other genome browsers.

### ChIA-View for visualization of ChIA-Drop in multiplex format

ChIA-DropBox introduces a new browser, named ChIA-View, specifically designed for visualizing single-molecule multiplex chromatin complexes. ChIA-View is implemented as an easy-to-use *R* Shiny app, and is pre-loaded with demo data and also allows users to upload their own data.

The input file for a ChIA-Vview is a “REGION” file, which is one of ChIA-DropBox’s standard output file formats with one chromatin fragment per line. The “REGION” file includes the fragment coordinates, the GEM ID (which includes the barcode), and the fragment count of the GEM. The “REGION” file may optionally contain an additional column that indicates whether each fragment indicates a gene promoter (“P”) or not (“N”).

ChIA-View generates displays of the putative chromatin complexes as follows. First, the user specifies the coordinates of a genomic region of interest, and then ChIA-View selects all fragments within the genomic region (*bedtools* v2.27.1). Next, a matrix is constructed in which each column is a bin along the genomic region and each row is a chromatin complex. For each chromatin complex (determined based on the GEM ID with the barcode), each column is assigned a value of 1 if that genomic bin contained a fragment for that chromatin complex (otherwise the column is assigned a value of 0).

Next, hierarchical clustering is applied to this Boolean matrix to arrange the different chromatin complexes by similarity along the y-axis. Specifically, the clustering is applied by first computing a similarity matrix based on the distances between all pairs of chromatin complexes and then selecting a linkage criterion for the clustering. These configurations are specified by the parameters “clustering_distance_rows” (= “euclidean” in default) and “clustering_method”(= “complete” in default) of *R* package *pheatmap* (v1.0.10) in default. Then, a heatmap of the binned and clustered ChIA-Drop complex can be displayed using the *R* package *pheatmap* (v1.0.10). At this point, ChIA-View is ready to display the ChIA-Drop data in “Cluster view” mode.

Once the clustering has been achieved, ChIA-View can also readily switch to displaying the data in “Fragment view”. In this display mode, the ordering of the chromatin complexes along the y-axis is preserved, but now each chromatin complex is displayed with its original fragment coordinates and a thin line connecting them all. The “Cluster view” and the “Fragment view” modes are well suited for unenriched ChIA-Drop data.

ChIA-View also offers two other display modes that are particularly well suited for promoter-centric ChIA-Drop libraries, such as RNAPII-enriched ChIA-Drop libraries. In the “Promoter view” mode, chromatin complexes that overlap gene promoters are filtered and retained (based on the additional annotation column in the “REGION” file). Then, chromatin complexes are arranged along the y-axis based on which gene in the region they overlap. Chromatin complexes are displayed such that promoter-involving fragments are displayed in a bright color and non-promoter-involving fragments are displayed in a dark color. In the “Gene body view”, the user selects a specific gene, then chromatin complexes involving the promoter of that gene are filtered and retained, and complexes are sorted along the y-axis based on the span of the chromatin complex.

Other *R* packages used in clustering heatmap view and linear fragment view include: *pheatmap* (v1.0.10), *ggplot2* (v3.0.0), *dplyr* (v0.7.6), *bedr* (v1.0.4), *ggforce* (v0.1.3), *cowplot* (v0.9.3), *gtable* (v0.2.0), *gridExtra* (v2.3), *reshape2* (v1.4.3), *RColorBrewer* (v1.1-2), *ggrepel* (v0.8.0), and *ggpubr* (v0.1.7).

